# The transcriptional landscape of cultivated strawberry (*Fragraia* × *ananassa*) and its diploid ancestor (*Fragraia* × *vesca*) during fruit development

**DOI:** 10.1101/2020.01.02.893453

**Authors:** Yongping Li, Tianjia Liu, Huifeng Luo, Shengcai Liu

## Abstract

Cultivated strawberry (*Fragaria ananassa*) comes from four diploid ancestors: *F.vesca, F.viridis, F. iinumae and F.nipponica*. Among them, the *F.vesca* is the most dominance subgenome for cultivated strawberry. It is not well understood how gene expression contributes to differences during fruit development between diploid and octoploid strawberry. Here, we used comprehensive transcriptomic analyses of *F.vesca* and *F*. × *ananassa* to investigate gene expression at different stages of fruit development. In total, we obtained a total of 3,187 (turning stage) and 3,061 (red stage) differentially expressed genes with the pairwise comparisons between diploid and octoploid. Genes involved in flavonoids and phenlypropanoids biosynthesis, were almost up-regulated in the both turning and red stages of octoploid, and we also discovery a ripe-fruit specific module associated with several flavonoids biosynthesis genes, including *FveMYB10, FveMYB9/11*, and *FveRAP* by using weighted gene coexpression network analysis (WGCNA). Furthermore, we identified the species-specific regulated network in the octoploid and diploid fruit. Notably, we found that the WAK and F-box genes were enriched in the octoploid and diploid fruits, respectively. As a whole, this study contributes to shed new light on the flavonoid biosynthetic and fruit size of strawberry, with important implications for future molecular breeding in the cultivated strawberry.

## Introduction

Whole-genome duplication (WGD) or polyploidy, is an important evolutionary force in seed plants and angiosperms ^1^, which probably contributed to the varied and novel phenotypes. Polyploids are generally grouped into two types, autopolyploidy and allopolyploids, which involves in single or different diploid progenitors, respectively ^2^. Allopolyploids originate primarily from interspecific hybridization and composed of the complicated genetic background, so it contains a range of genetic alterations ^3,4^. The composition of different subgenomes can significantly repress the phenotype of allopolyploids and commonly reflects evolutionary and domestication influences ^5^. There are numerous crop species are allopolyploids, such as sorghum, cotton, potato, wheat, and sugarcane. Cultivated strawberry (*Fragraia* × *ananassa*) is also an allopolyploid (octoploid) species, whose fruit exhibit large size, disease-resistant, intense color, varied aroma and flavor, compared with diploid varieties.

Transcriptional remodeling in polyploids is caused by the interaction of diverged parental genomes, that is reunited in the allopolyploid and genome duplication. It is likely to contribute to heterosis and exhibit variation in adaptation to new conditions ^5^. Polyploidy changes the progress of meiosis and mitosis and the relationship between cellular components, through epigenetic changes to modify gene expression ^5,6^. Furthermore, gene dosage balance, trans- and/or cis-regulatory network also affects the gene expression. In the process of the polyploidization, the splicing sites could be further altered with the changes in the component, abundance and activity of splicing factor, which could affect the absence and presence of alternative splicing (AS) in polyploids ^7^. The AS could create new isoforms and hence change the proteins, and also could regulate transcripts expression level through triggering nonsense-mediated decay (NMD) pathway degradation ^8^. The AS events of diploid and octoploid strawberry have been identified ^9,10^, but we do not know the difference between them, especially in the process of fruit development.

Strawberry fruit is known as a source of polyphenol compounds, especially flavonoids, a large family of secondary metabolites. The highest content of flavonoids is proanthocyanidins (PAs), followed by anthocyanins, flavonols and the other phenolics ^11,12^. Fruit color is one of the breeding objectives of great interest in strawberry. The anthocyanin biosynthetic genes or transport genes have been identified in strawberry ^12,13^. Down-regulation of strawberry *FaGT1* can redirection of flavonoid biosynthesis in ripening fruit ^14^. The *FaMYB10* plays a general regulatory role in anthocyanin biosynthesis during strawberry ripening15. Furthermore, there are some critical genes involved in anthocyanin biosynthesis or transport, such as *FaMYB9/FaMYB10* ^16^ and GST ^17^. The availability of genome and transcriptome provides an opportunity to reveal the flavonoid biosynthetic pathway. However, no large-scale omics analysis has been performed to dissect the molecular mechanism of flavonoid biosynthesis in strawberry. Flavonoids have potential health benefits and anthocyanins may be involved in plant growth and development, stress responses and gene expression.

The plant cell wall is mainly composed of cellulose, hemicellulose, pectin, and proteins. The wall was thought to be involved in modulating plant development ^18^. Cell expansion ^19^ and pathogen infection ^20^ could alter biosynthesis and cell wall components and downstream cytoplasmic events such as systemic acquired resistance. These events showed communication between the cytoplasm and the cell wall is of great importance. Wall-associated kinases (WAKs) are one of the four classes of proteins that physically link the ECM to the plasma membrane and may promote communication between the two compartments. WAKs belong to receptor-like kinases, which have a cytoplasmic protein kinase domain and linked to the pectin fraction of the plant cell wall. WAKs proteins regulate diverse biological processes, and main function is involved in the pathogen response and plant cell expansion. The numerous environmental stimuli can activate their expression.

*Fragaria vesca* is a wild diploid strawberry and subgenomic donor of the octoploid genome of cultivated strawberry (*F*. × *ananassa*) ^21^. However, the underlying mechanism and the ultimate consequences of subgenome dominance have been still mostly unknown. Here, we successfully collected the spatial and temporal transcriptome data of diploid and octoploid strawberry fruit tissues, at different stages of fruit ripening. Global transcriptome analysis showed that octoploid strawberry exhibited different transcriptome profiles in comparison with the diploid strawberry. It is helpful to reveal the transcriptional network and to identify potential genes and functional classes involved in strawberry fruit ripening. As a whole, this study contributes to shed new light on the flavonoid biosynthetic and fruit size of strawberry, with important implications for future molecular breeding in the cultivated strawberry.

## Materials and methods

### Plant materials

The receptacle was collected from the diploid *F. vesca* YW5AF7 ^22^ and a popular cultivar ‘Sweet Charlie’, which were germinated in a greenhouse under natural photoperiod. For both varieties, the receptacle fruits were collected in the morning at late March. The receptacle with achenes removed was quickly frozen in liquid nitrogen and stored at −80°C until RNA isolation.

### Illumina RNA-sequencing of pooled receptacle tissue from YW5AF7 and ‘Sweet Charlie’

Equal amount of receptacle tissues at small green to ripe stages were pooled for each variety (*F. vesca* YW5AF7 and *F*. × *ananassa* “sweet Charlie”). Total RNA was extracted and evaluated as described above. 6-16 μg of total RNA per sample was sent to Beijing Genomics Institute (BGI, Wuhan, China) for strand-specific library construction and sequencing on the platform of Illumina HiSeq2500. 89 and 142 million 125bp paired-end reads were generated for YW5AF7 and Sweet Charlie, respectively. The RNA-seq data were uploaded to the Sequence Read Archive (SRA) of the National Center for Biotechnology Information (NCBI) (http://www.ncbi.nlm.nih.gov/sra/), under the accession number SRX1756111, SRX1756112 respectively.

### Description of previous diploid and octoploid fruit datasets

We collected total 80 Illumina-based datasets representing 36 different fruit tissues of diploid and octoploid fruit at different developmental stages, including seven development stages for diploid receptacle (cortex and pith) and other fruit tissues (wall, style, embryo, ovule, ghost) ^23,24^. as well as the four development stages (green, white, turning, red) for octoploid achene and receptacle ^25^.

### Read mapping and transcript assembly

Raw data were quality- and adaptor-trimmed using Fastx-Tookit (http://hannonlab.cshl.edu/fastx_toolkit/). Next, clean reads were aligned to *F. vesca* Genome V4.0 genome ^26^ downloaded from GDR using the program STAR ^27^ with default parameters. Stringtie ^28^ was used to individually assembly transcripts based on a reference guided assembly strategy. Putative transcripts for each sample were merged into a unique set of transcripts by TACO software ^29^. The merged transcripts file was then compared with the v4.0.a2 ^30^ annotation GTF file using cuffcompare ^31^.

### Differential expression analysis

The gene-level count matrices for annotation genes were generated by FeatureCount ^32^. The median-of-ratios method was used to normalize for RNA composition and sequencing depth using DESeq2 package ^33^. DESeq2 software was used to identify differentially expressed genes (DEGs) based on the negative binomial generalized linear model. The DEGs were determined based on an absolute log2-fold change value of >4 and *padj* < 0.001.

### Functional classification of genes based on MapMan pathway and GO enrichment

The MapMan software ^34^ was employed to facilitate the assignment of DEGs into functional categories (bins). MapMan bins of *F.vesca* V4 genome annotation were assigned by the Mercator pipeline (https://mapman.gabipd.org/app/mercator). GO terms of the full set of genes in v4.0.a2 annotation were assigned by using Blast2GO ^35^, and significantly enriched GO terms were identified based on a hypergeometric test with false discovery rate (FDR) < 0.05.

### AS event and Differential AS gene detection

To classify the AS events, the rMATS software ^36^ was employed with the transcripts GTF files assembled from diploid and octoploid data. Five major types of AS events, namely retained intron (RI), alternative 3’ splice site (A3SS), alternative 5’ splice site (A5SS), skipped exon (SE) and mutually exclusive exons (MXE), were counted for each sample, respectively. rMATS was also used to detect the differentially spliced events between the diploid and octoploid, using the aligned BAM files as input with default parameters. The merged transcripts GTF file generated by cuffcompare software was used as the reference. AS event between diploid and octoploid with an FDR < 0.05 and ‘Exon Inclusion Level Difference’ > 0.1 were defined as the differentially spliced events.

### Validation of Differential AS events by RT-PCR

Total RNA was isolated from the pooled receptacle of diploid and octoploid using the Plant Total RNA Isolation Kit (Sangon Biotech, Shanghai, China, No. SK8631). The cDNA was synthesized using a PrimeScript RT reagent kit with gDNA Eraser (TaKaRa); 5x diluted cDNA served as the template. KOD DNA polymerase (TOYOBO, Bio-Tech, Cat# F0934K) was used for PCR amplification. Finally, the PCR products were visualized in agarose gel stained.

### Weighted gene co-expression network analysis

Co-expression network analysis was performed using the WGCNA package in R ^37^. The raw counts for 26,194 genes (TPM > 2 in at least one tissue) were normalized to homoscedastic expression values using *varianceStabilizingTransformation* function in DESeq2 package ^33^. A signed hybrid network was being constructed using the automatic network construction function blockwiseModules calculated the topographical overlap matrices (TOM) with default settings, except that the power = 20, TOMType = “signed”, minModuleSize = 30 and mergeCutHeight = 0.25. To determine the association of module with the tissue-specific expression for each tissue, the cor () function was used for the correlations correlation between modules and tissues. A summary profile (module eigengene, ME) was calculated for each module via PCA. The most highly connected genes are central hub genes. Finally, the network was visualized using Cytoscape_v3.6.1 ^38^.

## Results

### Comparative analysis of transcriptome sequencing data between the diploid and octoploid strawberry

YW5AF7 (*F. vesca*) is a diploid wild strawberry mainly used for basic research, whereas *F*. × *ananassa* is an octoploid commercial variety. The fruit flesh of *F*. × *ananassa* is quite different from YW5AF7 in terms of size, color, aroma, and texture (Fig. S1A). To understand the effect of ploidy on global gene expression, we collected a total of 80 RNA-seq libraries generated from 36 different tissue types, including four fruit development stages (green, white, turning, red) for octoploid achene and receptacle, as well as seven development stages for diploid receptacle (cortex and pith) and other fruit tissues (wall, style, embryo, ovule, ghost). In total, 2.3 billion RNA-seq reads were generated from diploid and octoploid receptacles. As *F. vesca* is a near-complete genome and well annotation genes, contributing to one set of the subgenomes of *F*. × *ananassa*, its genome was used as a reference in our analyses. The clean reads were mapped to the *Fragaria vesca* V4 genome 26 with annotation ver4.0.a2 ^30^, with a mapping ratio of 95% for diploid samples and 85% for octoploid samples, respectively (Table S1). The correlation between each tissue is in Figure 1. The transcriptome profile from octoploid exhibited a large distance from diploid strawberry. The clustering results indicated that the polyploid effect could largely explain the variation among samples. Overall, our results suggested that the quantity and quality RNA-seq data were suitable for downstream comparative analyses.

**Figure 1.**
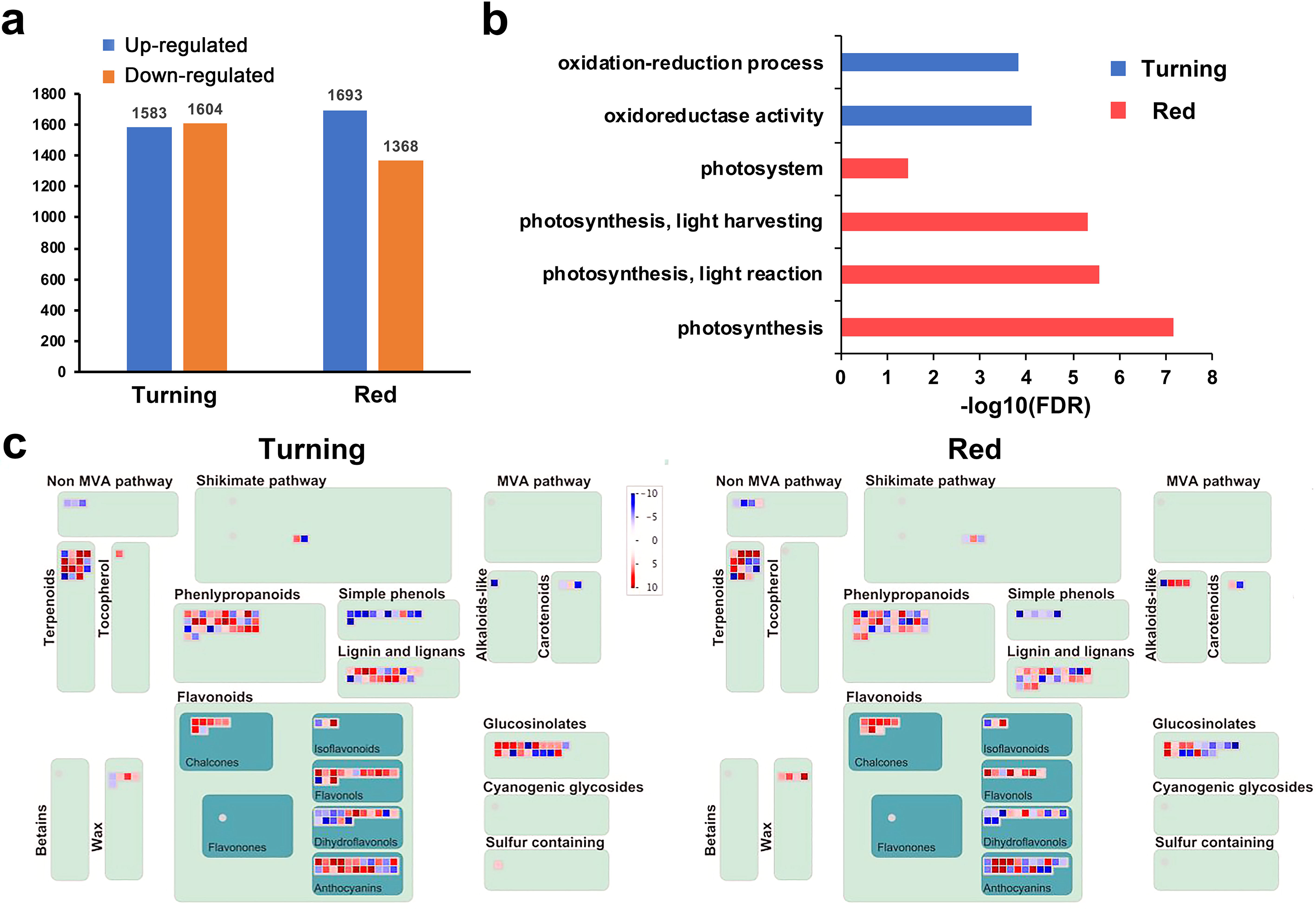
Summary of genes differentially expressed (DEG) between diploid and octoploid in turning and red stage. a. Numbers of detected DEGs between diploid and octoploid in turning and red stage. b. The gene ontology (GO) enrichment of the up-regulated genes during the turning and red stage. c. Secondary metabolism overview corresponding to the DEGs in the turning and red stage. The color key indicated the log2(fold change) between diploid and octoploid.

### Transcriptome altered between diploid and octoploid during fruit ripening

Turning and red are two representative stages of strawberry fruit ripening. The RG15d and RG22d samples were turning and red stages in diploid, consistent with turning receptacle and red receptacle in octoploid, respectively. To gain a genome-wide view of the gene expression at the two stages, we collected the RNA-seq datasets of the turning and red stages for both octoploid and diploid strawberry fruit (Figure S1). Then, pairwise comparisons (RG15d vs. turning receptacle, RG22d vs. red receptacle) were performed to identify differentially expressed genes using DESeq2. We obtained a total of 3,187 (turning) and 3,061 (red) differentially expressed genes between diploid and octoploid. At turning stage, pairwise comparisons of octoploid and diploid showed that 1,583 genes were significantly upregulated, while 1,604 were significantly downregulated (|fold change| > 4,*padj* ≤ 0.001) (Table S2). At red stage, 1,693 and 1,368 genes were significantly upregulated and downregulated, respectively, in samples of the red stages octoploid compared with diploid. Further analysis indicated that only three genes were up-regulated in the two sets of pairwise comparisons, including two pectin lyase-like superfamily protein genes (FvH4_6g41380 and FvH4_1g03310), and an *AtCAX1* homologous gene, FvH4_6g11493, which was involved in cellular homeostasis and oxidative stress in Arabidopsis ^39^. Whereas only one gene (FvH4_4g11920) was down-regulation.

To further understand the potential functions of differentially expressed genes between octoploid and diploid, we carried out gene ontology (GO) enrichment analysis of the up-regulated genes in octoploid samples. Interestingly, GO analysis showed that “oxidation-reduction process” and “oxidoreductase activity” terms (oxidation-reduction related) were enriched in turning stage; “photosystem”, “photosynthesis, light harvesting”, “photosynthesis, light reaction” and “photosynthesis” terms (photosystem related) were enriched in red stage (Figure 1b).

Furthermore, MapMan bins were also used to identify specific categories of genes or pathways which were enriched in differentially expressed genes (DEGs). As shown in Figure 1c, the secondary metabolism pathway analysis suggested that the genes involved in flavonoids and phenlypropanoids, especially chalcones, were almost up-regulated in both turning and red stages of octoploid.

### AS events between diploid and octoploid receptacles

To investigate the impacts of ploidy on AS and the contributions of AS to fruit quality, fruit (receptacle) transcriptomes from *F*. × *ananassa* cultivar ‘Sweet Charlie’ and *F. vesca* were obtained by Illumina RNA-seq. After comparing the isoforms in diploid and octoploid, a greater number of isoforms was uniquely identified in octoploid (28,662 octoploid versus 17,759 diploid) despite enough sequencing depth in our data (Figure 2a). The AS events in diploid and octoploid were further classified into different types, including RI, A3SS, A5SS, SE and MXE by the rMATs tools. Consistent with isoforms number, more AS events were also identified in octoploid (Figure 2b). For both varieties, RI is the most frequent AS type, followed by SE, A3SS, A5SS and MXE, similar to other plant species ^40–42^. The result indicated that the global AS regulation in octoploid is similar to that in diploid.

**Figure 2.**
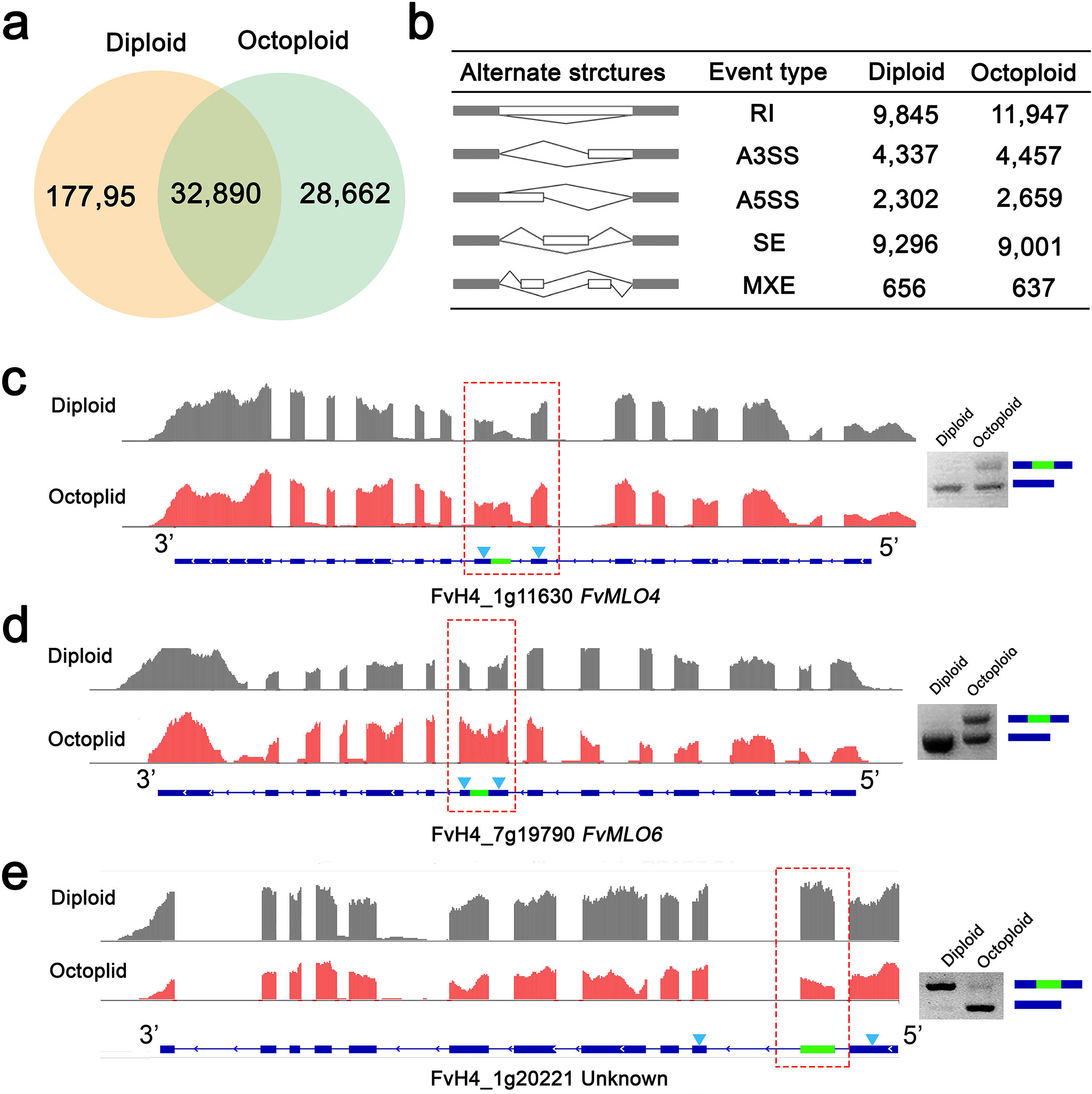
Large-scale identification of alternative splicing in diploid and octoploid by RNA-seq. a. Venn diagram showing the common and unique transcripts from diploid and octoploid. b. Statistics of different AS events obtained from the diploid and octoploid. c,d,e. PCR validation of the differential alternative splicing events of *FvMLO4, FvMLO6* and FvH4_1g20221 between diploid and octoploid. For each gene, read coverage graphs viewed in IGV were shown with the red dotted square indicating the differentially spliced regions between the two samples; gene model is shown on the bottom with blue arrowheads indicating the position of primers used for RT-PCR validation; agarose gel image shows the bands of two transcript variants amplified from the same RNA samples for sequencing with gene models on the right side.

It is difficult to determine the expression level of each isoform obtained by Illumina RNA-seq. AS events were individually compared between the diploid and octoploid by using rMATS to identify potential AS genes that regulated fruit quality ^36^. As shown in Table S3, 1,779 and 2,284 AS events, respectively, were more abundant in octoploid and in diploid (FDR < 0.05, absolute “Exon Inclusion Level Difference” > 10%). For example, FvH4_1g11630, named *FvMLO4* encoding a homolog of the barley mildew resistance locus o (MLO) protein ^43^, has one more 3’ splicing site in the eighth exon with a 41.9% higher exon inclusion level in octoploid which results in a premature stop codon and is validated by RT-PCR (Fig. 2c). Interestingly, another MLO protein-encoding gene FvH4_7g19790, called *FvMLO6* ^43^, also possesses a differential splicing event with a retained intron (Fig. 2d). It suggests that the MLO gene family is susceptible to the AS regulation and potentially contributes to the powdery mildew resistance of Sweet Charlie. Besides, FvH4_1g20221, encoding an unknown protein, also exists in *Malus domestica* and *Prunus persica*, whose isoform skipped the entire second exon specifically in octoploid leading to delete 71 amino acids (Fig. 2e). In summary, although the differences in AS events were found in the receptacle diploid and octoploid strawberries, the characteristics of AS regulation are similar.

### Coexpression network analysis with WGCNA

The dynamic expression patterns of genes reflect their roles during fruit development. RNA-seq data for 36 different tissue/stages were generated from diploid (28 samples) and octoploid (8 samples). To comprehensively understand the gene regulatory network of the two varieties, and to identify specific genes associated with polyploids and flavonoids biosynthetic pathway during fruit ripe, we performed the weighted gene coexpression network analysis (WGCNA). After filtering the low expression genes (TPM > 2 in at least one tissue), 26,194 genes were retained for WGCNA. The cluster dendrogram revealed the correlation of transcriptomes of different tissues in diploid and octoploid (Fig. 3a), after using the ‘hclust’ function matrix. Different stages of the same tissue from diploid and octoploid are similar and could be clustered together.

**Figure 3.**
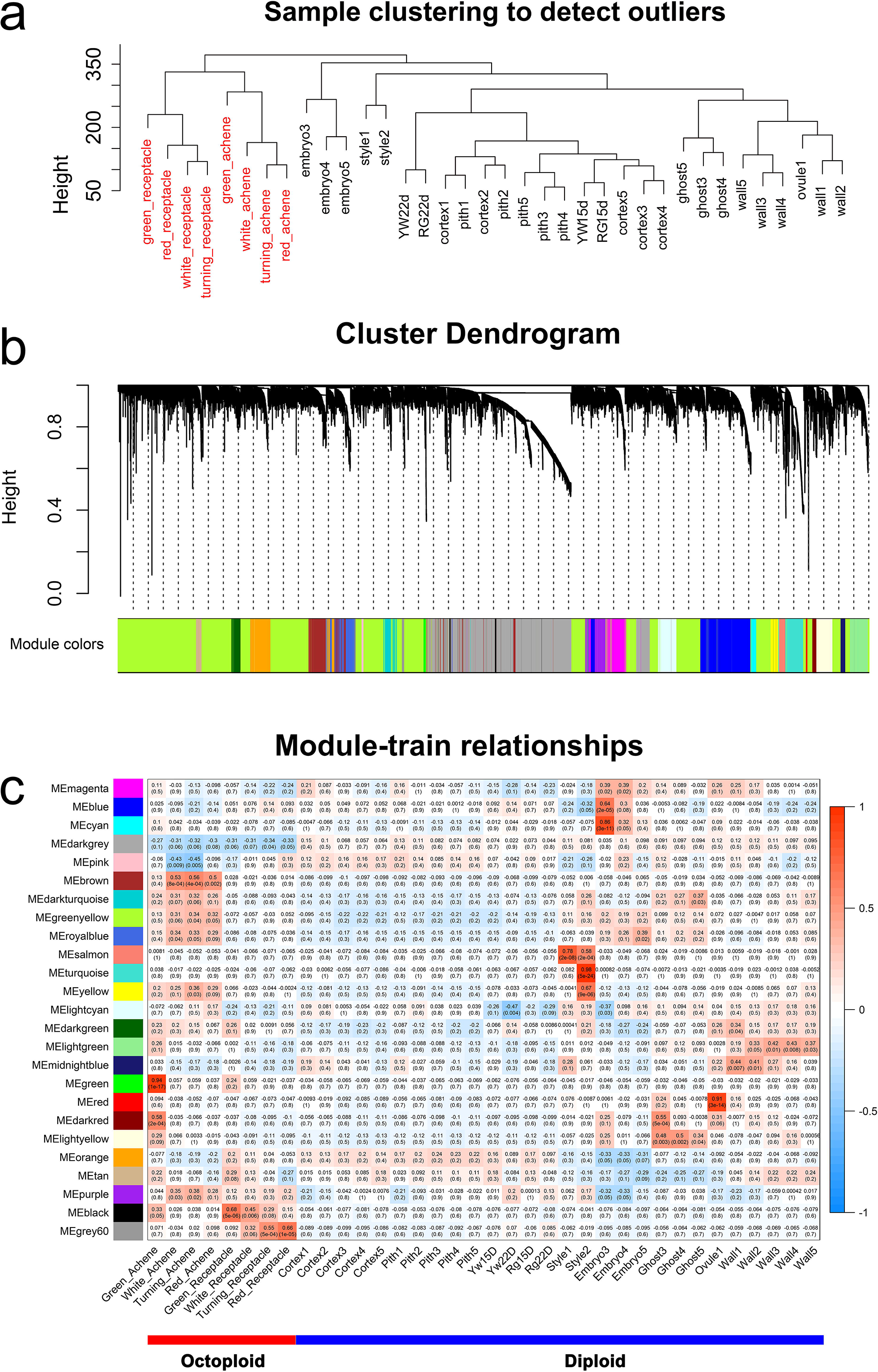
Coexpression network analyses of flower, fruit, and vegetative tissues. a. Cluster dendrogram showing global relationship between different samples. The y axis is the degree of variance. b. Dendrogram showing co-expression modules identified by WGCNA. Each leaf in the tree is one gene. The major tree branches constitute 26 modules labeled by different colors. c. Heatmap showing cluster-tissue associations of the network. Each row corresponds to a module. Each column corresponds to a specific tissue/stage. The color in the boxes indicates the correlation coefficient between the module and the tissue type.

Coexpression networks were constructed on the basis of pairwise correlations of gene expression of all sample tissues. Using standard parameters, the genes could be assigned to 26 distinct modules (labeled by different colors) (Fig. 3b). Consistent with cluster analysis, the tissues were a major driver of module identity, according to eigengenes tissue-specific expression profiles (Fig. 3c, Table S4), which allows for visualization of the cluster-tissue association. For instance, genes in salmon color module are the highest expression in style1. Brown color module correlates with octoploid achenes. A large number of tissues and stages of the diploid and octoploid enabled the development of fruit coexpression networks.

### Receptacle-Associated modules provide insight into metabolism regulation network during fruit ripening

The most obvious phenomenon during fruit ripening is the synthesis of anthocyanins. Previous transcriptome analysis revealed that flavonoids biosynthesis genes were induced transcription in the receptacle, implicating the importance of these genes ^24^. The grey60 color module (Table S6) is correlated with the octoploid receptacle development, and the correlation is getting higher as the receptacle matures (Figure 3c). Figure 4a showed the eigengene expression of the grey60 module. A heatmap showed the relative NRPKM of genes from the receptacle module and revealed that many genes were high expression both in octoploid ripening fruit (Figure 4b) and in diploid (Yw22D and Rg22D). This result indicates that this module is highly associated with fruit ripening, whether in diploid or octoploid. WGCNA is also employed to construct gene networks, and the networks were visualized using Cytoscape (Figure 4c). Two MYB genes (FvH4_3g1399 and FvH4_3g13990) were found among genes. FvH4_1g22020 (*FvMYB10*) could regulate the flavonoid/phenylpropanoid metabolism during strawberry fruit ripening ^15^. FvH4_3g13990, is a homolog of Arabidopsis *AtMYB67* (AT3G12720) gene, which is a member of the R2R3-MYB factor gene family. The gene ontology (GO) terms (Biological Process) in grey60 color module are “lipid metabolic process”, “transferase activity”, “carbohydrate binding” and “catalytic activity” (Figure 4d, Table S5). KEGG of metabolism-related pathways were significantly enriched among the grey60 genes, such as ‘Metabolism’, ‘metabolism of terpenoids and polyketides’ and ‘pyrimidine metabolism’ (Figure 4e).

**Figure 4.**
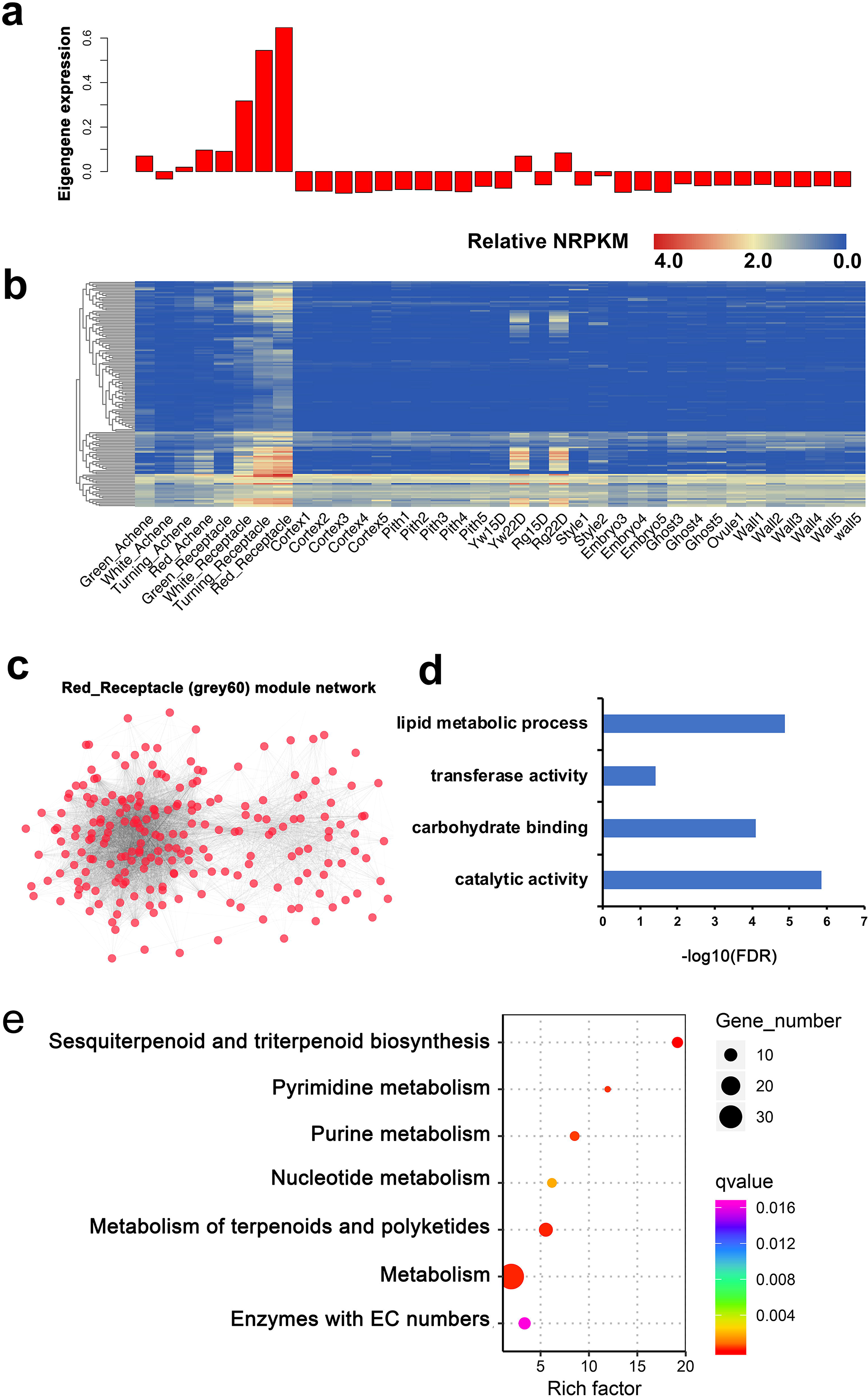
Ripe fruit associated genes and networks. a. Eigengene expression profile for the ripe-fruit module in different tissues. The y axis indicates the value of the module eigengene; the x axis indicates the sample type. b. The correlation network of ripe-fruit module. Each colored circle (node) represents one gene. c. The correlation network of ripe-fruit (grey60) module. d. GO terms are enriched in the ripe-fruit module genes. e. The enrichment analysis of KEGG pathway for the genes in ripe-fruit module. The area of each colored circle is proportional to the number of genes involved in each pathway, the color indicates the *q*-value, and the *x*-axis is the Rich Factor.

### Species-specific modules and hub genes were identified through WGCNA

Of particular interest is the identification of the species-specific gene modules due to there are many differences in the octoploid and diploid fruits. To identify conserved genes across species, we used the samples from all stages of fruit development and performed WGCNA analysis. Notably, 6 out of 26 coexpression modules (darkgrey, brown, green, purple, black and grey60; p-value < 0.001) were composed of genes that were highly expressed in a single species (Figure S2). Among them, the darkgrey and black color modules were the most significant module for each species. Figure 5a, b, showed the eigengene expression for the darkgrey (diploid) module and black (octoploid) module, respectively.

**Figure 5.**
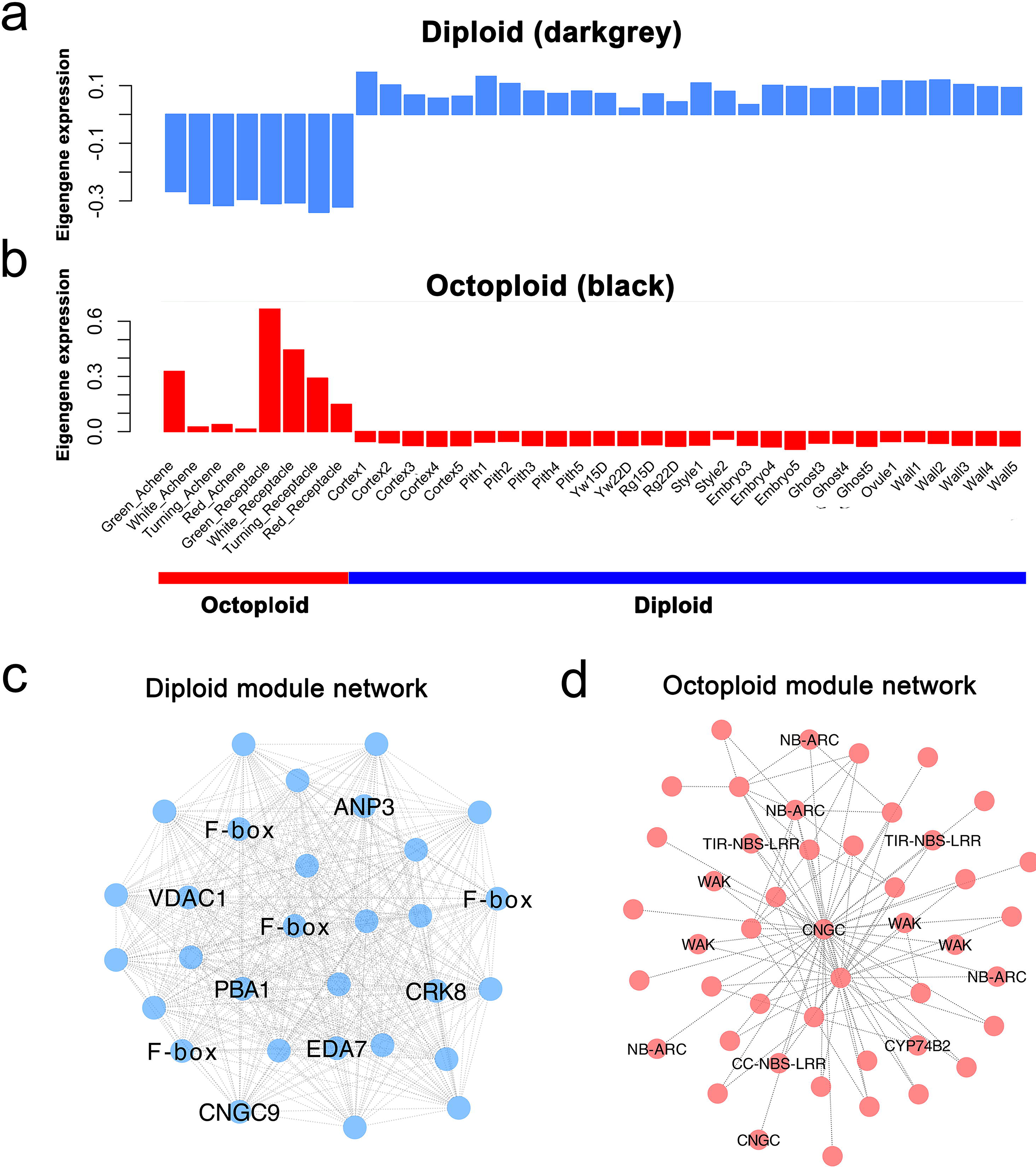
Diploid and octoploid-specific genes and networks. a, Eigengene expression profile for the darkgrey (diploid) module in different tissues. b. Eigengene expression profile for the black (octoploid) module in different tissues. The y axis indicates the value of the module eigengene; the x axis indicates sample type. c. The correlation network of darkgrey (diploid) module. d. The correlation network of black (octoploid) module. Genes with the edge weight higher than 0.2 are visualized by Cytoscape.

The darkgrey (diploid) module (Fig. 5a, Table S7) is a positive correlation with diploid fruit tissues and a negative correlation with octoploid fruit tissues. To detect the ability of the consensus clusters to predict functional relationships between genes, we performed the GO enrichment analysis. The GO terms (Biological Process) in diploid (darkgrey) module were related to “cell communication”, “signal transduction” and “celluar process” et al (Table S5). We also used WGCNA ^37^ to construct gene networks, in which each node represents a gene and the connecting lines (edges) between genes represent coexpression correlations, WGCNA identifies the most highly connected, or the most central genes within each module, referred to as “hub genes”. The top 10 genes in each of the specific modules are shown in Table 1.

**Table 1.**
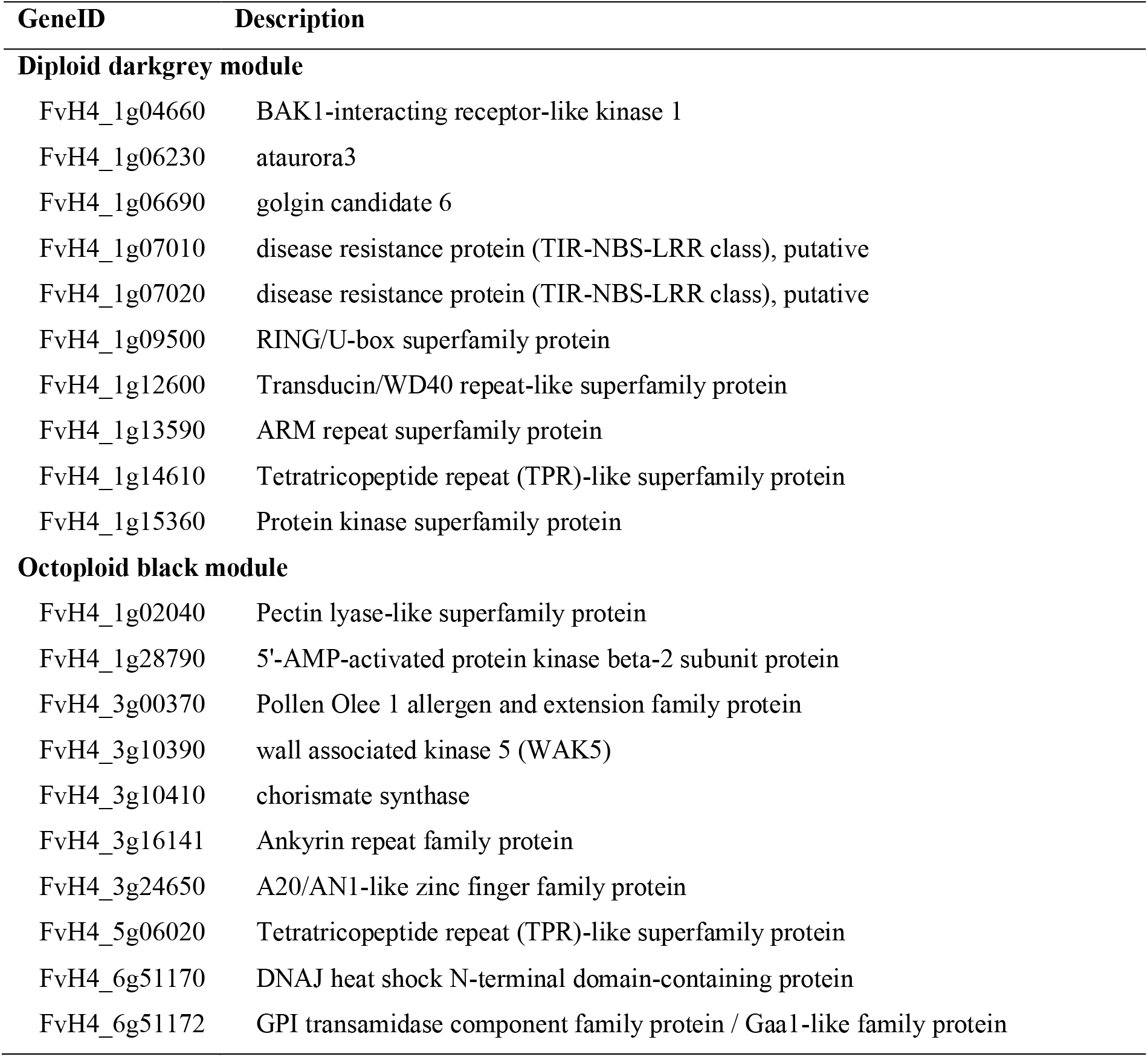
Candidate hub genes (top10) in diploid and octoploid modules, respectively.

In the darkgrey (diploid) module network (Fig. 5c; Table S7), many of the network genes are F-box gene. Interesting, the previous study showed that the F-box genes and miRNA formation the miRNA-FBX networks in *F. vesca* fruit and FBX-targeting miRFBXs might have evolved to regulate *F.vesca-specific* processes ^44^. Besides, the hub genes in diploid module include BAK1 (FvH4_1g04660), ataurora3 (FvH4_1g06230) and two TIR-NBS-LRR (FvH4_1g07010 and FvH4_1g07020) et al (Table 1).

In contrast, the black (octoploid) module is a positive correlation with octoploid fruit tissues (Figure 5b, Table S8), and a negative correlation with diploid fruit tissues. The GO terms (Biological Process) in black module are related to “cellular protein modification process” (Table S5). Strikingly, many wall associated kinases (WAKs) genes werein the octoploid (darkgrey) module network (Figure 5d). The top 10 hub genes list including one WAK gene, which might be involved in cell wall-related processes (Table 1).

## Discussion

### A large number of genes exhibit differentially regulated AS events between the diploid *F. vesca* and octoploid *F*. × *ananassa*

The genus *Fragaria* is comprised of both cultivated (*F*. × *ananassa*) and wild species with various ploidy levels, such as diploid, tetraploid, hexaploid, octoploid and decaploid ^45^. Thus, it is an ideal genus for ploidy research. *F. vesca* is the progenitor diploid species of *F*. × *ananassa*, and their fruit quality differs in terms of size, color, aroma, and texture. Therefore, comparing the transcriptome landscapes in receptacle between *F. vesca* and *F*. × *ananassa* could help gain insights into the regulation of both ploidy and fruit quality. Interesting alterations of AS in some genes have been observed in *Brassica* and watermelon between diploid and tetraploid varieties ^7,46^. However, these papers only examined a limited number of genes that would unlikely reflect the change of the entire transcriptome. In this study, the overall features of AS are the same between diploid and octoploid plants, for instance, the percentage of each major AS type is similar (Figure 2), indicating that the splicing machinery works the same. As the *F*. × *ananassa* species originates from natural hybridization between *F. virginiana* and *F. chiloensis*, and thus the *F*. × *ananassa* cultivar Sweet Charlie has overcome the transcriptome shock appeared in the first few generations of allopolyploidy or autopolyploidy ^47,48^. We cannot infer the AS features during the transcriptome shock.

In summary, a large number of genes exhibit differentially regulated AS events between the diploid *F. vesca* and octoploid *F*. × *ananassa*, but the general AS characteristics are comparable between these two strawberries with different ploidy levels; Altogether, this study identified the majority of alternatively spliced events during receptacle development in strawberry, serving as a valuable resource for future functional studies.

### Flavonoids biosynthesis pathway during fruit ripening

The strawberry fruit ripening contains two-phase flavonoid formation stages. The first stage is the formation of flavanols and the second is related to anthocyanin and flavonol accumulation ^49^. Strawberry ripe fruits contain high concentrations of flavonoids, especially the red colord anthocyanins ^12^. Before fruit ripening, the anthocyanins are mainly present in the form of PAs. Subsequently, it will decrease considerably during fruit ripening ^11,16^.

In this study, the standard networks were generated by the WGCNA analysis pipeline. Our network analyses revealed five flavonoid biosynthesis related genes (*FvGST, FvMYB10, FvMYB9/11, FvUFGT, FvH4_1g02120*) in ripe-fruit module (Figure 4), which prompted us to investigate flavonoids biosynthesis pathway during the ripening of fruit. In strawberry, the *FaMYB9/FaMYB11, FabHLH3* and *FaTTG1* are homolog to Arabidopsis *AtTT2, AtTT8* and *AtTTG1*, which form a ternary protein complex to regulate the expression of the PA biosynthetic genes ^16^. *MYB10*, a R2RMYB transcription factor, is repressed expression mainly in the fruit receptacle. However, it is activated by abscisic acid (ABA) and repressed by auxins in the process of the fruit ripening. It plays an important regulatory role in the flavonoid/phenylpropanoid pathway during the ripening of strawberry ^15^. A reduced anthocyanin in petioles (rap) mutant, was identified to be a premature stop codon in a glutathione S-transferase (GST) *FvGST* gene. *FvGST* mediates foliage and fruit pigmentation and also acts downstream of the fruit-specific *FvMYB10* ^17^. In Arabidopsis, the homologs of strawberry *FvUFGT* encodes an anthocyanidin 3-O-glucosyltransferase which involved in the biosynthesis of flavonoid glucosides. The FvH4_1g02120, which homologs of *Arabidopsis AT1G49390* gene was also involved in the flavonoid biosynthetic process. Overall, the synthesis of anthocyanins in strawberry fruit involves a very complex regulatory network.

### Wall-associated kinases (WAKs) may function in pathogen response and cell expansion of octoploid strawberry

It is an interesting finding that many WAK genes were found in octoploid module network (Figure 5d). The wall-associated kinases (WAKs), are receptor-like kinases, which have a cytoplasmic protein kinase domain and linked to the pectin fraction of the plant cell wall. The plant cell wall is mainly composed of cellulose, hemicellulose, pectin, and proteins. The wall was thought to be involved in modulating plant development (Kohorn, 2000). The cell expansion (Cosgrove, 1997) and pathogen infection (Hammond-Kosack and Jones, 1996) could alter biosynthesis and cell wall components and downstream cytoplasmic events such as systemic acquired resistance. These events showed communication between the cytoplasm and the cell wall is very important. Wall-associated kinases (WAKs) are one of the four classes of proteins that physically link the ECM to the plasma membrane and may promote communication between the two compartments. WAKs belong to receptor-like kinases, which have a cytoplasmic protein kinase domain and linked to the pectin fraction of the plant cell wall. We all know, cultivated strawberry (*F*. × *ananassa*) is an allopolyploid (octoploid) species, which fruit exhibit large size, disease-resistant, intense color, varied aroma and flavor, compared with diploid varieties. It is in accord with many WAK genes that existed in octoploid module network. WAKs proteins regulate diverse biological processes.

FBX proteins regulate diverse biological processes and were grouped into two general types based on whether they function in conserved processes, such as embryogenesis and circadian rhythms, or relatively specialized processes, such as pollen recognition and pathogen response.

## Acknowledgment

This work was supported by the Program for High-level University Construction of the Fujian Agriculture and Forestry University (612014028) and the Natural Science Foundation of Fujian Province (2018J01700).

## Author contributions

Conceived and designed the experiments: YL and SL. Analyzed the data: YL. Performed the experiments: TL, HL. Wrote the paper: YL and SL.

## Supplementary data

Figure S1. Photographs of diploid and octoploid fruit tissues in turning and red stages.

Figure S2. Heat map showing cluster-tissue association of the WGCNA network.

Table S1. Summary of RNA-seq read statistics

Table S2. DEGs between diploid and octoploid in turning and red stages.

Table S3. The differential alternative splicing events in fruit receptacles between diploid and octoploid.

Table S4. Data for WGCNA analysis.

Table S5. WGCNA modules gene list and GO terms in grey60, black and darkgrey module.

Table S6. Grey60 module network file.

Table S7. Darkgrey module network file.

Table S8. Black module network file.

---

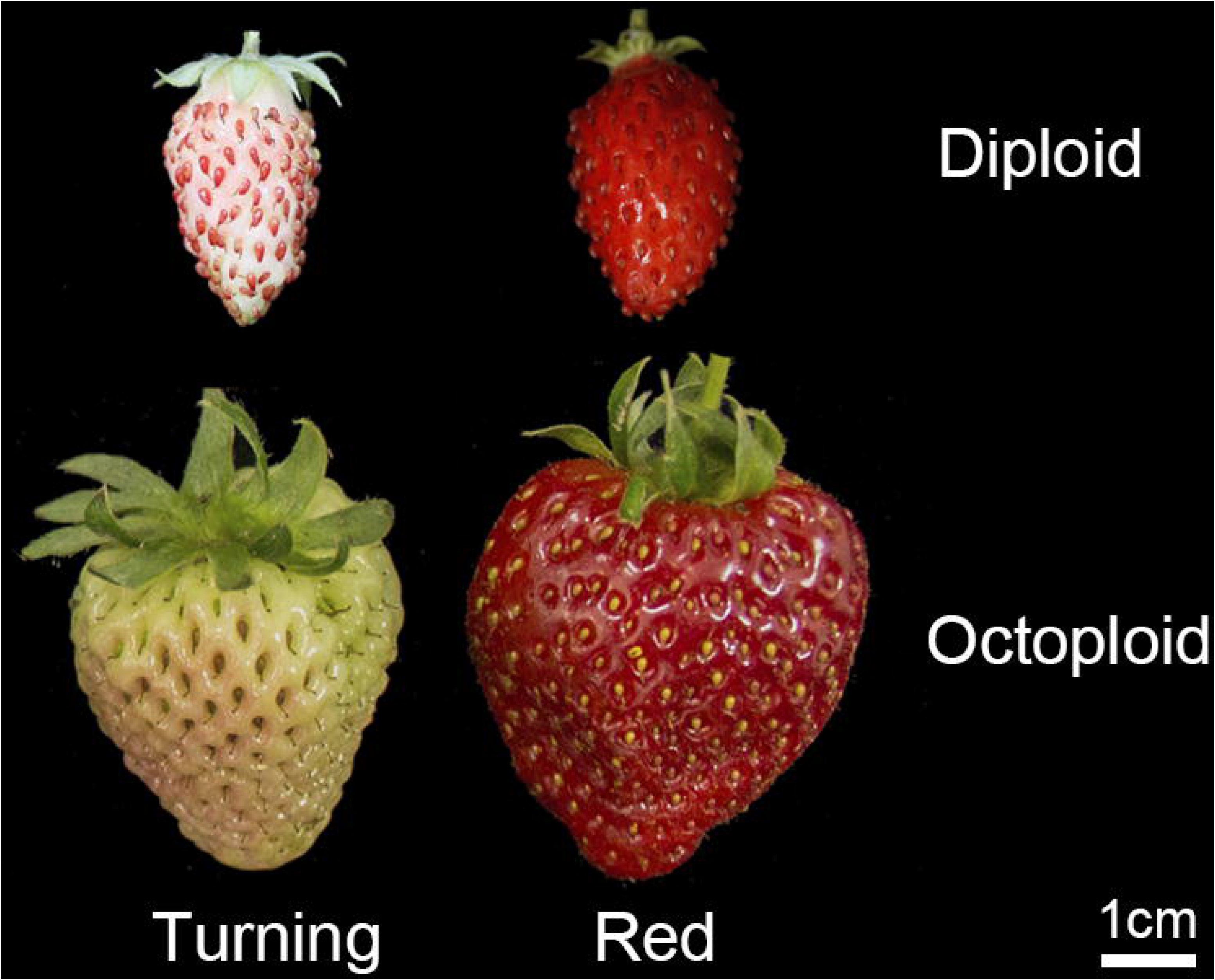

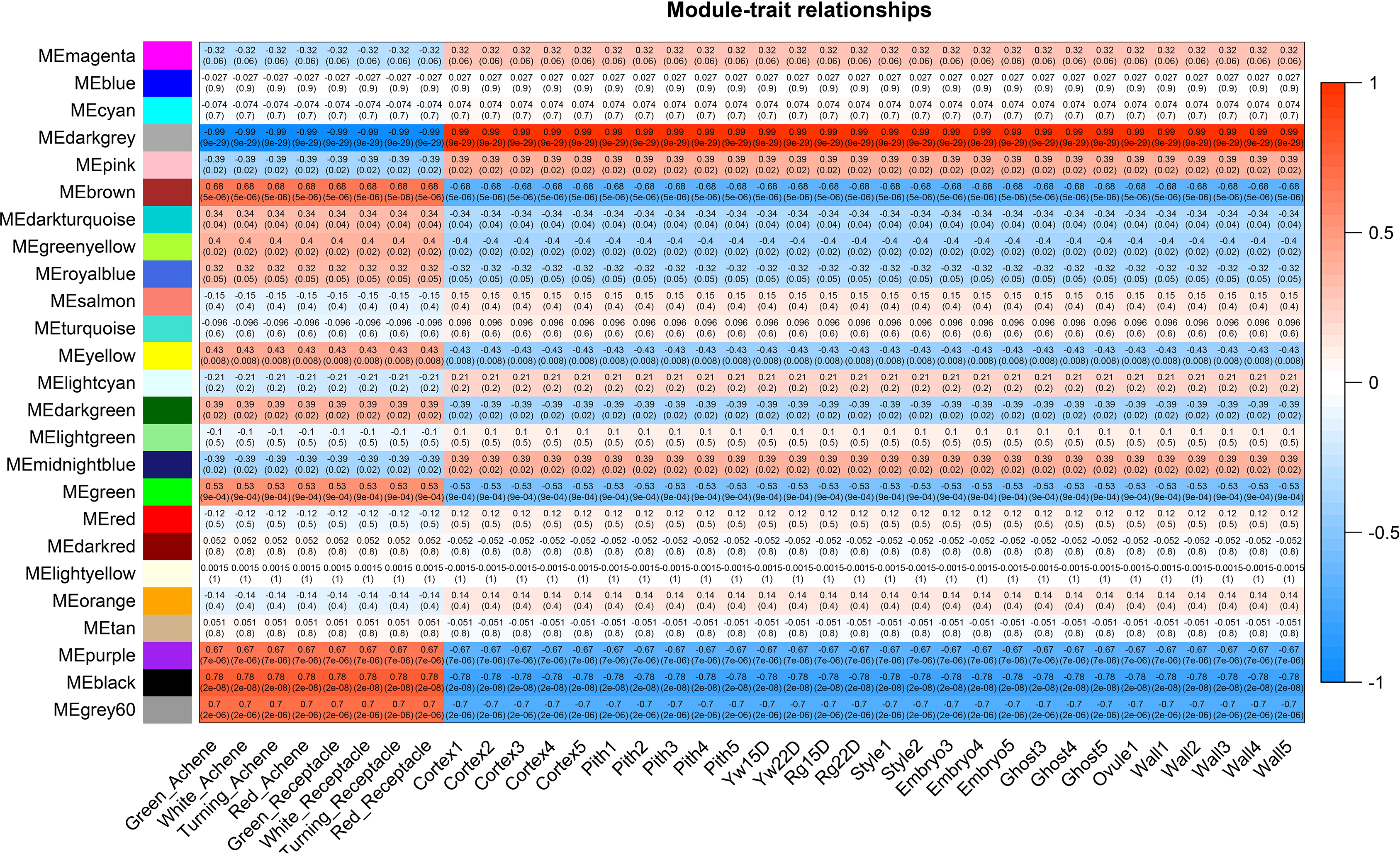

